# QuiXoT: quantification and statistics of high-throughput proteomics by stable isotope labelling

**DOI:** 10.1101/193607

**Authors:** Marco Trevisan-Herraz, Inmaculada Jorge, Elena Bonzon-Kulichenko, Pedro Navarro, Jesús Vázquez

## Abstract

In most software tools for quantification of mass spectrometry-based proteomics by stable isotope labelling (SIL), there is a recurrent disconnection between the use of a statistical model and convenient data visualisation to check correct data modelling. Most of them lack a robust statistical framework, using models which do not account for the major difficulties in proteomics, such as the unbalanced peptide-protein distribution, undersampling, or the correct separation of sources of variance. This makes especially difficult the interpretation of quantitative proteomics experiments. Here we present QuiXoT, an extensively tested quantification and statistics open source software based on a robust and extensively validated statistical model, the WSPP (*weighted spectrum, peptide, and protein*). Its associated software package allows the user to visually represent and inspect results at all modelled levels (scan, peptide and protein) on routine bases. It is applicable to practically any SIL method (SILAC, iTRAQ, and ^18^O among others) or MS instrument.

## 1 Introduction

Quantitative high-throughput proteomics experiments require accurate, robust and powerful statistics to allow reliable data analysis. One of the main challenges in the field is the lack of tools to inspect the accuracy of the quantitative results. Here we present the QuiXoT software package, a whole software platform developed in C# that allows quantification, thorough statistics using the WSPP statistical framework (Navarro, 2014), and graphical data inspection of any kind of high-throughput SIL experiment on routine basis.

QuiXoT has been extensively and successfully used in numerous works, demonstrating the robustness and adaptability of the algorithm (Jorge, 2009; Bonzon-Kulichenko, 2011a; Ramírez–Boo, 2011; Burillo, 2013; Perez-Hernandez, 2013; Jorge, 2014; Navarro, 2014; Burillo, 2015; García-Marqués, 2016; Zenón, 2016).

## 2 Main features

**Fig. 1.**
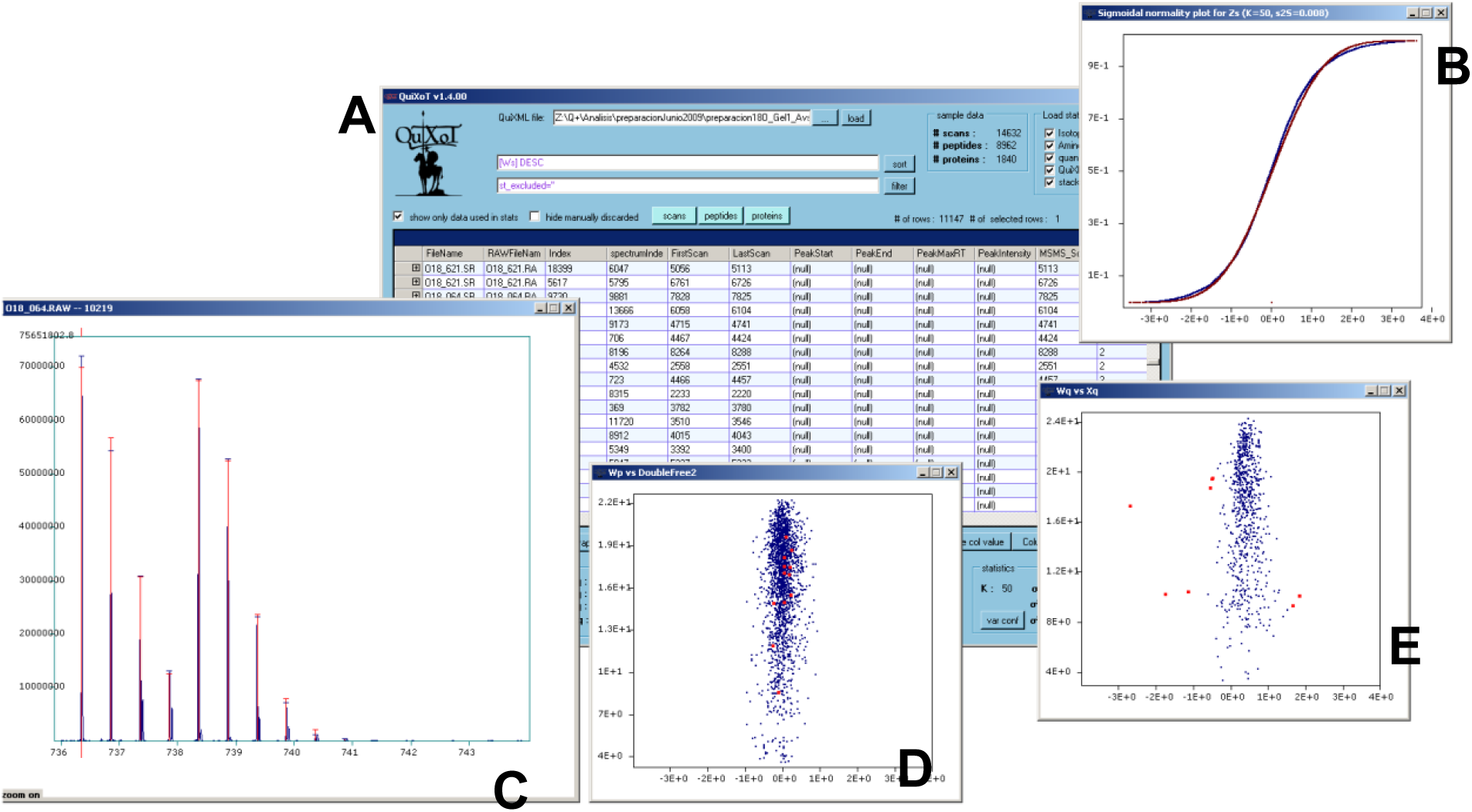
A screenshot of QuiXoT, showing some of the different graphs used during an analysis including: the main window, including the datagrid with information about the identification and quantification of spectra, peptides and proteins (A); a sigmoidal, cumulative normality plot, showing the experimental curve in blue vs the theoretical N(0,1) distribution in red (B); the quantification of an ^18^O experiment, comparing the experimental spectrum in blue, and the theoretical spectrum in red (C); centred peptide fold-change vs peptide weight, highlighting in red peptides from the same protein (D); protein fold-change vs protein weight, highlighting in red proteins with statistically significant expression changes (E).

### 2.1 Quantification

QuiXoT is able to quantify practically any kind of isotopic or isobaric labelling, such as different types of SILAC, ^18^O, iTRAQ or TMT, and it has been optimised to work with several instruments of different resolution capabilities, ranging from LTQ XL to Orbitrap Fusion instruments (Thermo), or TOF-based spectrometres like the 4800 Proteomics Analyzer (AB SCIEX). The software is able to perform a retention time alignment of labelled and unlabelled peaks for the cases the chromatography system (typically UPLC) separates slightly in elution time labelled peptides from their unlabelled pairs. This avoids quantification biases and missing features. QuiXoT quantification can only be made for Thermo’s RAW files, but tools for importing quantifications from other sources are available (see Supplementary Information).

In MS1 quantitative SIL methods, such as SILAC or ^18^O, QuiXoT uses the total intensity area of peptide isotopic profiles, and distributes it in labelled and unlabelled intensity areas according to a Newton-Gauss algorithm. This is especially relevant for controlling the labelling efficiency of each peptide for labellings, which have more than one labelling state (i.e. ^18^O labelled peptides may have one or two ^18^O atoms exchanged at the carboxy-terminus).

The goodness of each quantification is evaluated by using the experimental peptide signal, and it is used as the statistical weight at scan level, needed for the WSPP model (Jorge, 2009).

### 2.2 WSPP statistical model

As mentioned, QuiXoT can apply, after the quantification is finished, the WSPP (*weighted scan, peptide, protein*) statistical model (Navarro, 2014), which resolves most of the major difficulties in the analysis of quantitative proteomics experiments. Using this model, the experiment error is separated into scan, peptide, and protein error sources, allowing to compare the results obtained by different SIL approaches. Moreover, this hierarchical error model is advantageous for localising the source of error at which experimental problems might arise and for optimising sample preparation protocols and MS instrument settings; for instance, an increased scan-level variance shows a problem in the mass spectrometer, whereas an alteration of the peptide-level variance reveals potential flaws in the digestion, or other steps during peptide handling. The model has been intensively validated by confronting numerous data distributions versus different null-hypothesis experiments, and its performance has proven to be at least similar to other well-known models, which are specific to each SIL method and mass spectrometre (Navarro, 2014).

Researchers that want to use their own quantification algorithms and resources can still use QuiXoT for the statistical analysis, by parsing tables in text file to the QuiXoT format, using a dedicated tool for it (see Supplementary Information).

### 2.3 Graphic tools

To help the analysis and interpretation, QuiXoT comes with a flexible graphical interface that provides 2D scatter graphs of any two numerical columns in the main datagrid, such as expression change (in 2-based log scale) vs statistical weight, as well as useful graphs to check labelling efficiency and normality of experimental data. Plots are dynamically connected to the datagrid, so that the use of filters or a zoom in the plot will cause the datagrid to display affected elements (scans, peptides, or proteins): selecting rows in the datagrid highlights the corresponding points in the plot. This allows the user to quickly identify outliers, expression changes, and biases (Bonzon-Kulichenko, 2011b); see how estimated quantification values fit with their corresponding spectra and visualise on the 2D plots the scans of peptides coming from a protein of interest, etc. Users can also see how estimated quantification values fit with their corresponding spectra.

## 3 Workflow

**Fig. 2.**
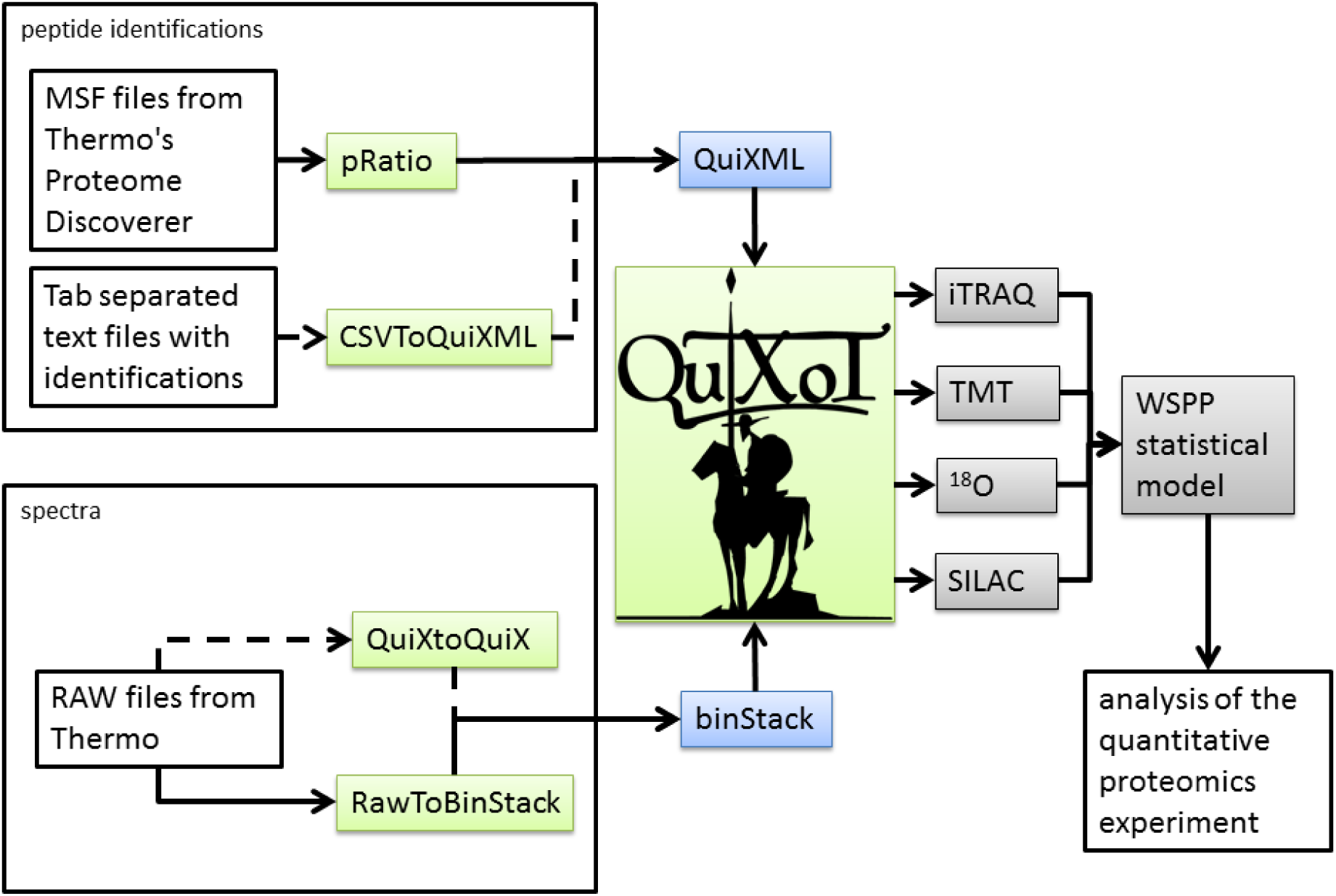
The workflow used by QuiXoT. Two elements are used as general input: an XML file (QuiXML) used as a master document and a folder containing spectral information in a set of binary files, which is used to quantify over spectra (called binStack). The QuiXML file is the output of pRatio (a software to validate peptide identifications from Proteome Discoverer), but alternatively identifications from any other software are possible as far as they are presented in a tab-separated text file, using the CSVToQuiXML tool. Spectra from Thermo RAW files can be processed using RawToBinStack to generate the binStack folder; additionally, MS1 spectra can be analysed and processed using the QuiXtoQuiX software. QuiXoT is optimised to be used for experiments quantified using iTRAQ, TMT, ^18^O, or SILAC. Once spectra are quantified, the WSPP model can be used to perform the statistical analysis. More information is available in the wiki.

QuiXML files are typically generated as an output file of pRatio, the software used to validate identifications using SEQUEST’s XCorr parametre (Eng, 1994). Alternatively, they can also be produced by parsing information from any source, stored as a tab-separated-value text file.

Additionally, there are a number of configuration files to allow full control of the software without changing the code. The most important being the QuantitationMethods.xml, an XML file containing different parametres for each SIL strategy. Information about this and other configuration files is available in the Supplementary Information, as well as in the abovementioned wiki. Programmers can also modify the source code, which is written in Visual C#, and publicly available under the Apache 2.0 licence.

The QuiXML file may be updated by the user with quantitative and statistical information from the WSPP statistical model. Parametres such as labelling efficiency (for ^18^O labelling), sample normality, or expression changes by low FDR can be checked here. There is as well the option of exporting the datagrid to a tab-separated-values text file.

The main workflow has been optimised to start from Thermo Proteome Discoverer’s MSF files, after validating its identifications using pRatio, whose output includes a QuiXML file. Spectra can be imported by storing them in a binStack folder from Thermo RAW files by using the RAWToBinStack satellite program.

Alternatives to the main workflow, such as importing already quantified data from other applications, can be found in the Supplementary Information.

The output can be stored in a tab separated text file, which can in turn be used as input for other statistical or visualisation software.

There is additional, extensive information about the management of the programs, workflows, DataGrid information stored in QuiXML files, unit tests, and analysis examples in the project’s wiki linked after the summary.

## 4 Examples

The whole process from identifications, protein quantifications and statistical analysis can be followed by downloading sample experiments available in the wiki. For a quick view, in the Supplementary Information the basic steps of the process are shown. For more details, the wiki includes the unit tests for four sample experiments, as well as a comprehensive analysis and practical examples. These experiments manage data using different combinations of three SIL techniques (SILAC, ^18^O and iTRAQ), two search engines (SEQUEST and Mascot) and three instruments (LTQ Orbitrap XL, Orbitrap Elite, and MALDI TOF/TOF).

## 5 Conclusions

QuiXoT is a helpful tool for researchers who want to get the most out of their high-throughput quantitative proteomics experiments, allowing the practical use of the WSPP statistical model. Furthermore, its output can be used to perform systems biology analysis in connection with other computational resources available.

## 6 Funding

This work was supported by grants BIO2012-37926 and BIO2015-67580-P from the Spanish Ministry of Economy and Competitiveness and grant PRB2 (IPT13/0001 - ISCIII-SGEFI/FEDER, ProteoRed) and RD12/0042/0056 from RIC-Red Temática de Investigación Cooperativa en Salud (RETICS), Fondo de Investigaciones Sanitarias, Instituto de Salud Carlos III. The CNIC is supported by the Spanish Ministry of Economy and Competitiveness (MINECO) and the Pro-CNIC Foundation, and is a Severo Ochoa Center of Excellence (MINECO award SEV-2015-0505).

## 7 Contribution of each author

PN, MTH, and JV designed QuiXoT; PN and MTH developed the software; IJ and EBK benchmarked QuiXoT and introduced critical input; MTH and PN wrote the manuscript.

